# Molecular Mechanism of Huanglian Jiedu Decoction in Treatment of Alzheimer’s Disease based on Network Pharmacology

**DOI:** 10.1101/2024.05.15.594364

**Authors:** Qiuyan Ye, Xue Li, Wei Gao, Yutong Zhang, Miao-miao Zhang, Liping Zheng, Honglin Li

## Abstract

**Purpose:** The aim of this study was to investigate the pharmacological mechanism of Huanglian Jiedu Decoction (HLJDD) in treating Alzheimer’s disease through network pharmacology. HLJDD is a classic Chinese medicine prescription that is recommended in the Chinese Alzheimer’s disease diagnosis and treatment guidelines. However, the mechanism of HLJDD treatment for Alzheimer’s disease (AD) remains unclear because of its complicated components.

**Methods:** The related ingredients and targets of HLJDD in treating AD were screened by Traditional Chinese Medicine Systems Pharmacology Database and Analysis platform (TCMSP), TTD, OMIM, GeneCards, and DrugBank. The data of the protein-protein interaction (PPI) network was constructed using STRING. The Metascape database was adopted for Gene Ontology (GO) functional annotation and Kyoto Encyclopedia of Genes and Genomes (KEGG) enrichment analysis. AutoDockTools was used for molecular docking verification.

**Results:** In the treatment of AD, HLJDD demonstrated strong efficacy with its core active components including were quercetin, β-sitosterol, stigmasterol, targeting key proteins such as AKT1, TNF, and IL6. Molecular docking tests confirmed the significant binding affinity between these components and the aforementioned targets. The biological pathway of HLJDD in treating AD primarily involves the modulation of IL-17, TNF, and other inflammatory cytokines to regulate their impact on nerve functions.

**Conclusions:** HLJDD may treat AD by inhibiting neuroinflammation through a comprehensive, multi-component, multi-target, and multi-pathway method.

## Introduction

Alzheimer’s disease (AD) is the most common type of senile dementia and is characterized by an insidious and progressive degenerative process of the central nervous system. Memory dysfunction, thinking, and behavioral issues are the primary clinical features. The pathogenesis of AD is highly complex and remains inconclusive, with mainstream hypotheses including the Aβ cascade reaction, Tau protein abnormality hypothesis, cholinergic hypothesis, neuroinflammation, metabolic disorders, and vascular injury [1]. Currently, FDA-approved clinical drugs for AD primarily consist of acetylcholinesterase inhibitors (AchE) and N-methyl-D-aspartate (NMDA) receptor antagonists. However, these drugs target a single factor, exhibit low drug resistance, and often lead to adverse effects, resulting in unsatisfactory clinical outcomes for AD treatment. Chinese medicine compound therapy has the advantages of multiple targets, multiple components and multiple pathways of action. Therefore, traditional Chinese medicine possesses unique benefits for the treatment of AD.

Huanglian Jiedu decoction (HLJDD) was initially documented in *The Handbook of Prescriptions for Emergencies*. It consists of four herbs: Coptidis rhizome (Huanglian or HL), Scutellariae radix (Huangqin or HQ), Phellodendri chinensis cortex (Huangbo or HB), and Gardeniae fructus (Zhizi or ZZ) in Chinese. HL serves as the sovereign medical, while HQ and HB, the minister medicinals, are responsible for clearing the heat of triple energizers. ZZ acts as both assistant medicinal and courier medicinal, aiding in clearing the heat of triple energizers and directing heat downward. Several studies have revealed the primary active components of HLJDD, including cognitive improvement [2], anti-inflammatory [3], anti-endotoxin [4], anti-atherosclerosis [5], and other effects. HLJDD is recognized as a classical heat-clearing medicinal agent and is widely employed in treating moderate and severe dementia because of its beneficial effects.

Network pharmacology is a comprehensive discipline that integrates the theories of systems biology, pharmacology, information networks and computer science. Using computer simulation and various databases, network pharmacology screens drug molecular targets and disease-related targets. By employing network visualization and network analysis systems, it sheds light on the complex network connections among drugs, targets, and diseases. It constructs a “drug-target” system to analyze and predict the action mechanism of drugs at multiple levels. The systematic and holistic features of network pharmacology align with the principle of holistic thinking emphasized by traditional Chinese medicine [6].

This study employs network pharmacological methods to determine the molecular mechanism of HLJDD in treating AD, because of its complex composition and multi-level effects. HLJDD shows multi-target effects in addressing AD, however the specific active components and mechanism of action remain unclear. The goal is to establish a theoretical foundation for future investigations. The workflow of the study is illustrated in **Figure 1**.

## Materials and Methods

### Collection and screening the active components of HLJDD

The components and targets of the four herbs in HLJDD were obtained from Traditional Chinese Medicine Systems Pharmacology Database and Analysis platform (TCMSP, https://old.tcmsp-e.com/tcmsp.php) [7]. Following TCMSP recommendations, the components were screened based on 2 thresholds of ADME: Oral bioavailability (OB) ≥30% and Drug-likeness (DL) ≥0.18. At the same time, the published literature was reviewed and organized to supplement the known targets of the undetected active components. The Uniprot database (https://www.uniprot.org/uniprot/) was used to translate the targets of the active components into gene symbols.

### Searching for potential AD-related targets of HLJDD

We conducted a search using “Alzheimer’s disease” as keyword to identify *Homo sapiens* genes associated with Alzheimer’s disease. AD-related targets for the treatment of AD were extracted from the TTD (https://db.idrblab.net/ttd/) [8], OMIM (http://www.omim.org/), GeneCards (https://www.genecards.org/) and DrugBank (https://go.drugbank.com/) databases. After consolidating all targets and removing duplicate entries, we compiled a list of targets related to AD by Venny (https://bioinfogp.cnb.csic.es/tools/venny/).

### Constructing the protein-protein interaction (PPI) network

We constructed the PPI by the STRING database (https://string-db.org) [9]. The biological species selected was *Homo sapiens (human)*, with the minimum interaction threshold set to “highest confidence” (>0.9).

### Topological analysis

Cytoscape3.9.1 was employed to build the network diagram encompassing the components,targets and pathways. Subsequently, CytoHubba, an embedded tool in Cytoscape, was used to analyze the network topology parameters of potential components. Screening was performed by Degree Centrality (DC) to determine the hub genes.

### Gene Ontology (GO) Functional Annotation and Kyoto Encyclopedia of Genes and Genomes (KEGG) Pathway Analysis

Input potential targets into Metascape, setting the threshold at P < 0.01 and selecting *Homo sapiens* as the biological species for both GO enrichment analysis and KEGG pathway analysis. Saved the data results and visualized them using bubble charts, mapped with micro-biosis (https://cn.bing.com).

### Molecular Docking Validation

Molecular docking is a method used to evaluate the ability of an active drug ingredient to bind to a target. The PDB database (https://www.rcsb.org/) was used to obtain the structural formulae of small molecules in PDB format. The docking of receptor proteins and small ligand molecules was performed using AutoDockTools, and the docking results were visualized through Pymol.

## Results

### 1. Active components of HLJDD

A total of 58 chemical components were identified through TCMSP active components screening, meeting the criteria of ADME (OB≥30%, DL≥0.18). Specifically, these components encompassed 11 species of HL, 28 species of HQ, 24 species of HB, and 9 species of ZZ. Notably, there were overlaps in the components, with 4 types common to both HL and HB, 1 shared among HL, HB, and ZZ, 2 present in both HL and HQ, and 2 shared across HQ, HB, and ZZ (**Table 1**). The analysis further revealed 128 targets for HL, 72 targets for HQ, 122 targets for HB, and 120 targets for ZZ. Upon consolidation and elimination of duplicates, a total of 138 unique targets were identified.

### 2. Prediction of potential anti-AD targets of HLJDD

A total of 2462 Alzheimer’s disease (AD)-related targets were obtained by combining data from the GeneCards database (targets with a score > 10) with information from the OMIM, TTD, and DrugBank databases. Duplicate targets were removed through a merging process. Subsequently, 137 overlapping targets were identified as potential targets for Huli Jingdan Decoction (HLJDD) in AD treatment. Consequently, a final set of 137 potential targets for HLJDD in the context of AD treatment was established (**Fig. 2**).

### 3. PPI network construction

The potential targets were analyzed using STRING to construct the PPI network (**Fig. 3**). The network contained 137 nodes and 337 edges, with an average node degree of 4.92 and an average local clustering coefficient of 0.511. Therefore, it can be seen from Figure.2 that the targets of HLJDD are closely related to each other in the treatment of AD, indicating that HLJDD may exert its efficacy by acting on multiple targets.

### 4. The “Components-Targets-Pathways” network diagram construction

CytoScape 3.9.1 was used to construct the network diagram of the “Components-Targets-Pathways” (**Fig. 4**), in which the top 10 active components (ranked by degree) were quercetin (MOL000098, A1), β-sitosterol (MOL000358, B1), stigmasterol (MOL000449, B2), kaempferol (MOL000422, ZZ1), wogonin (MOL000173, HQ2), isocorypalmine (MOL000790, HB2), palmatine (MOL000785, A3), (S)-Canadine (MOL001455, HB4), (R)-Canadine (MOL002903, HL1), coptisine (MOL001458, A5).

CytoHubba was then used for network topology analysis, with darker node colors indicating higher DC and more interaction. The results revealed top 10 core targets in AD treatment with HLJDD: TP53, AKT1, TNF, IL6, MAPK1, IL1B, ESR1, IFNG, FN1 andCCND1 (**Fig. 5**), with TP53 having a high DC of 29 (**Table 2**).

### 5. GO enrichment and KEGG Pathway analysis

The top 20 results obtained from the analysis using Metascape for the treatment of AD with HLJDD included the enrichment of the GO **(Fig. 6)** and KEGG pathways (**Fig. 7)**, respectively. The analysis revealed significant involvement in potential biological processes such as response to oxidative stress and reactive oxygen species. Moreover, molecular functions were identified, including neurotransmitter receptor activity and postsynaptic neurotransmitter receptor activity. The cellular components linked to these processes were found to be synapses, dendrites, and neuronal cell bodies. Additionally, the enriched pathways encompassed IL-17 signaling pathways and neurodegenerative pathways. Detailed results of the target pathway enrichment can be found in **Table 3**, providing a comprehensive overview of the findings.

### 6. Molecular docking validation

To analyze the top 5 active components and core targets, molecular docking was conducted using AutoDockTools 1.5.7 software. In this process, a lower binding energy indicates a more effective docking interaction between the component and the target. The docking results revealed that all binding energies were less than or equal to -4 KJ/mol, demonstrating superior binding between the components and targets (**Table 4**). The binding energies of the first 5 interactions were visualized using PyMol 2.2.0 software, as shown in (**Fig. 8**).

## Discussion

AD, as the most common form of dementia and a prominent neurodegenerative disease requiring attention, has imposed significant burdens on patients, their families, and society at large. The disease’s pathological mechanism is intricate, with previous studies highlighting the beneficial role of Chinese medicine compound prescriptions in its treatment [10, 11]. HLJDD, a renowned classic prescription, is known for its efficacy in clearing heat, detoxifying, and facilitating fire purging, specifically targeting dementia arising from phlegm, stasis, and heat. Baicalin can reduce neuroinflammation and oxidative stress in rats with Alzheimer’s disease by inhibiting microglial activation. It also regulates the levels of neurotransmitters such as methionine and glutamine, correcting imbalances and addressing energy metabolism disordersl [12, 13]. This was demonstrated by changes in peripheral system biomarkers SOD, GSH-Px, AMPK, SIRT-1, and MDA, indicating that HLJDD can improve cognitive function. Moreover, a 70% ethanol extract from gardenia can boost serum catalase activity, lower acetylcholinesterase levels, and improve rats’ spatial learning and memory [14]. These findings underscore the potential neuroprotective effects of HLJDD and its medicinal components in combating AD and enhancing cognitive function across various biological pathways.

In this study, the significant role of HLJDD in the treatment of AD was investigated through the employment of network pharmacology to screen its active components, which include quercetin, β-sitosterol, and stigmasterol. Quercetin, a compound found in HLJDD, has been shown to inhibit the neuroinflammatory process by downregulating pro-inflammatory cytokines. Additionally, quercetin has the ability to stimulate neuronal regeneration, exerting its neuroprotective role by antagonizing cytotoxicity induced by oxidative stress in neurons [15]. Beta-sitosterol has been demonstrated to decrease the deposition of Aβ in the brains of AD mice and partially restore synaptic function [16]. Furthermore, AMPK triggers the NF-κB and NLRP3 pathways. Stigmasterol inhibits the microglial inflammation response to Aβ42 oligomers and lessen neuroinflammation in AD mice [17]. Molecular docking results also suggested that active components such as quercetin, β-sitosterol, and stigmasterol exhibit strong binding with core targets, indicating their potential therapeutic efficacy in the treatment of AD within the context of HLJDD.

In this study, AKT1, TNF, and IL-6 were identified as the primary targets for AD treatment. AKT, a protein kinase, plays a crucial role in various biological processes and pathologies such as cell growth, proliferation, and neurodegenerative diseases [18]. Among the isoforms of AKT, AKT1 is highlighted, and elevated levels of AKT1 phosphorylation have been associated with Aβ deposition and Tau protopathy, impacting neuronal and synaptic damage [19]. Neuroinflammation, characterized by elevated levels of proinflammatory cytokines in the cerebrospinal fluid of AD patients, is closely linked to disease progression. TNF, comprising TNF-α and TNF-β subtypes, along with IL-6, contributes to this inflammatory response. Upon stimulation by Aβ, microglia are overactivated, leading to the release of pro-inflammatory cytokines like TNF-α and IL-6, exacerbating neuroinflammation and neurotoxicity in AD progression [20]. Pathway enrichment analysis revealed that HLJDD may influence AD treatment through its impact on IL-17, TNF, and other signaling pathways. The IL-17 signaling pathway, which promotes an inflammatory response, has been found to be associated with AD progression, affecting Aβ deposition, tau protein phosphorylation, and blood-brain barrier integrity in the early stages of AD. The involvement of IL-17 in cognitive dysfunction and neuroinflammation has been established [21, 22]. TNF, as a key mediator of neuroinflammation, can activate either neuroprotective or neurodegenerative cell responses through its receptors TNFR1 and TNFR2 [23]. Thus, HLJDD may have a therapeutic impact on AD by regulating IL-17, TNF, and other signaling pathways.

This study utilized network pharmacology and molecular docking as research methods to investigate the potential active components, targets, and pathways of HLJDD for treating AD. The research findings indicated that individual compounds in HLJDD could target multiple proteins, and the same protein could be involved in various signaling pathways. These results suggest that the therapeutic effects of HLJDD on AD may stem from the synergistic interactions among its multiple components, targets, and pathways. By integrating network pharmacology and molecular docking approaches, this study lays the groundwork for the clinical application of HLJDD in AD treatment, providing a preliminary scientific rationale. Moreover, it presents a promising avenue for further elucidating the specific mechanisms of HLJDD in treating AD and for exploring novel clinical strategies for managing the disease.

## Key Points

1. Huanglian Jiedu Decoction can alleviate Alzheimer’s disease’s symptoms.
2. Huanglian Jiedu Decoction can alleviate the pathological damage caused by Alzheim er’s disease through multiple ways and multiple targets.
3. It is necessary to further verify the mechanism of Huanglian Jiedu Decoction in tre ating Alzheimer’s disease.

## ACKNOWLEDGMENTS

This work was supported by the National Natural Science Foundation of China [82105 035].

## DISCLOSURES

The authors have nothing to disclose.

## Notes

### Competing Interest Statement

The authors have declared no competing interest.

### Summary of Updates

*Corresponding author. Second Clinical College of Heilongjiang University of Chinese Medicine, Harbin, China (Honglin Li)

